# Giant virus creates subcellular environment to overcome codon– tRNA mismatch

**DOI:** 10.1101/2024.10.07.616867

**Authors:** Ruixuan Zhang, Lotte Mayer, Hiroyuki Hikida, Yuichi Shichino, Mari Mito, Anouk Willemsen, Shintaro Iwasaki, Hiroyuki Ogata

## Abstract

Codon usage consonant with the cellular tRNA pool is important for efficient translation. However, many eukaryotic viruses, including amoeba-infecting mimiviruses, have codon usage that is largely deviated from that of their host, despite using the host machinery for translation. This raises the question of how these viruses cope with the mismatch between tRNA supply and demand. Here we show that Acanthamoeba castellanii mimivirus generates a subcellular area in the host cells to translate virus mRNAs. A combination of genome-wide ribosome profiling and RNA sequencing showed that ribosomes traversed along viral mRNAs with fewer pausing events than were observed on amoeba mRNAs. Frequently used codons in viral mRNAs had higher tRNA accessibility than the same type of codons in amoeba mRNAs. tRNA sequencing showed that the tRNA pool was not greatly altered during the infection even though the virus encodes tRNA genes. Instead, by *in situ* labelling, we found that viral mRNAs and newly synthesized proteins were localized at the periphery region of the viral factory, likely creating a unique environment to facilitate viral translation. Our data provide a perspective on how local translation assists the virus in overcoming the mismatch between tRNA supply and demand.

## Introduction

Codon usage in genes is unique in each genome. Even for synonymous codons that encode the same amino acid, the codon usage frequency varies. Preferentially used codons are associated with high tRNA copy numbers in fast-growing microorganisms such as *Escherichia coli*^1^. Recent studies highlighted the importance of synonymous codon usage in mRNA stability^2,3^. Therefore, the concordance of tRNA supply and demand is crucial for ensuring efficient gene expression.

Giant viruses are characterized by huge genomes (up to 2.5 Mb), a large number of genes (up to 2000 genes) for reprograming host cell metabolism, and large virions of over 1 µm, being comparable in size to some bacteria^4–7^. Infection by giant viruses induces a rapid shift in the host cell transcriptome, characterized by large occupancy of viral mRNAs^8–10^. However, the codon usage of some giant viruses is poorly adapted to the tRNA pool of the host^11,12^. For example, Acanthamoeba castellanii mimivirus (APMV) — the first virus isolate of family *Mimiviridae* — has an AT-rich genome (G+C content 28%), whereas the genome of its host *Acanthamoeba castellanii* is GC-rich (G+C content 58%)^13,14^. Consequently, the APMV codon usage varies extensively from the codon usage of *A. castellanii*^11,13^.

This raises the question of whether translation of APMV mRNAs is negatively impacted by the codon usage bias or whether APMV alleviates the unfavourable translation condition during infection. The genomes of *Mimiviridae* viruses encode many translation-related genes, including translation factors, tRNAs, and aminoacyl-tRNA synthetases, which were previously thought to be unique to cellular genomes^4–7^. Ribosomes localize around the viral factory (VF) of mimiviruses where viral DNA replication and transcription are carried out^15,16^. These observations implies that mimiviruses manipulate the translation system in infected host cells to overcome the codon usage mismatch.

To characterize the translation landscape of mimiviruses, we monitored changes in the transcriptome, tRNAs, and translatome during viral infection by RNA sequencing (RNA-Seq), optimized tRNA-Seq^17^, and ribosome profiling (Ribo-Seq)^18,19^. Our results indicate that the overall composition of the global tRNA pool was not significantly altered during APMV infection, with limited contribution of tRNAs encoded in the viral genome. At the early infection stage, the translation efficiency of the viral mRNAs was comparable to that of the host mRNAs despite using rare codons. Even at the later infection stages, the rare codons did not cause the ribosome to pause on viral mRNAs; rather, the accessibility to tRNAs for codons ending with A or U was different between viral and host mRNAs. The microscopic analysis using fluorescence *in situ* hybridization (FISH) and fluorescent non-canonical amino acid tagging (FUNCAT)^19^ showed that the spatial organization within infected cells indicated co-localization of viral mRNAs, host rRNAs, and newly synthesized proteins around the VF. Our results suggest that APMV created a subcellular heterogeneity for efficient translation of the viral genes.

## Results

### Imbalance of codon usage between APMV and the amoeba host

We characterized the dissimilarity in codon usage between APMV and the host *A. castellanii* (hereafter referred to as amoeba). Consistent with previous genome analyses^13,14^, the host and viral genes had distinct G+C content (Extended Data Fig. 1A) and codon usage (Extended Data Fig. 1B). APMV mRNAs predominantly use AU-rich codons, such as AAA, AAU, and AUU, whereas AU-rich codons were much less common in amoeba mRNAs. The tRNA gene copy number of amoeba was positively correlated with the amoeba codon usage (Extended Data Fig. 1C), whereas it showed no such correlation with the APMV codon usage (Extended Data Fig. 1D).

### Impact of APMV infection on RNA abundance and translation

To investigate the dynamics of transcription and translation of viral and host genes, we performed RNA-Seq and Ribo-Seq in APMV-infected amoeba cells at 0, 2, 4, and 8 hours post-infection (hpi) (Fig. 1A). Consistent with an earlier report^20^, the ribosomal footprints (hereafter footprints) showed two peaks in length at 21–22 and 29–30 nucleotides (nt) (Extended Data Fig. 2A, B). The footprint counts along genes showed clear 3-nt periodicity from the start codon (Extended Data Fig. 2C, D). Our data had high reproducibility for read mapping for both the host and viral genes (Extended Data Fig. 2E–H). These features indicate the robust detection of footprints generated by ribosomes translating mRNAs for both host and virus.

**Fig. 1.**
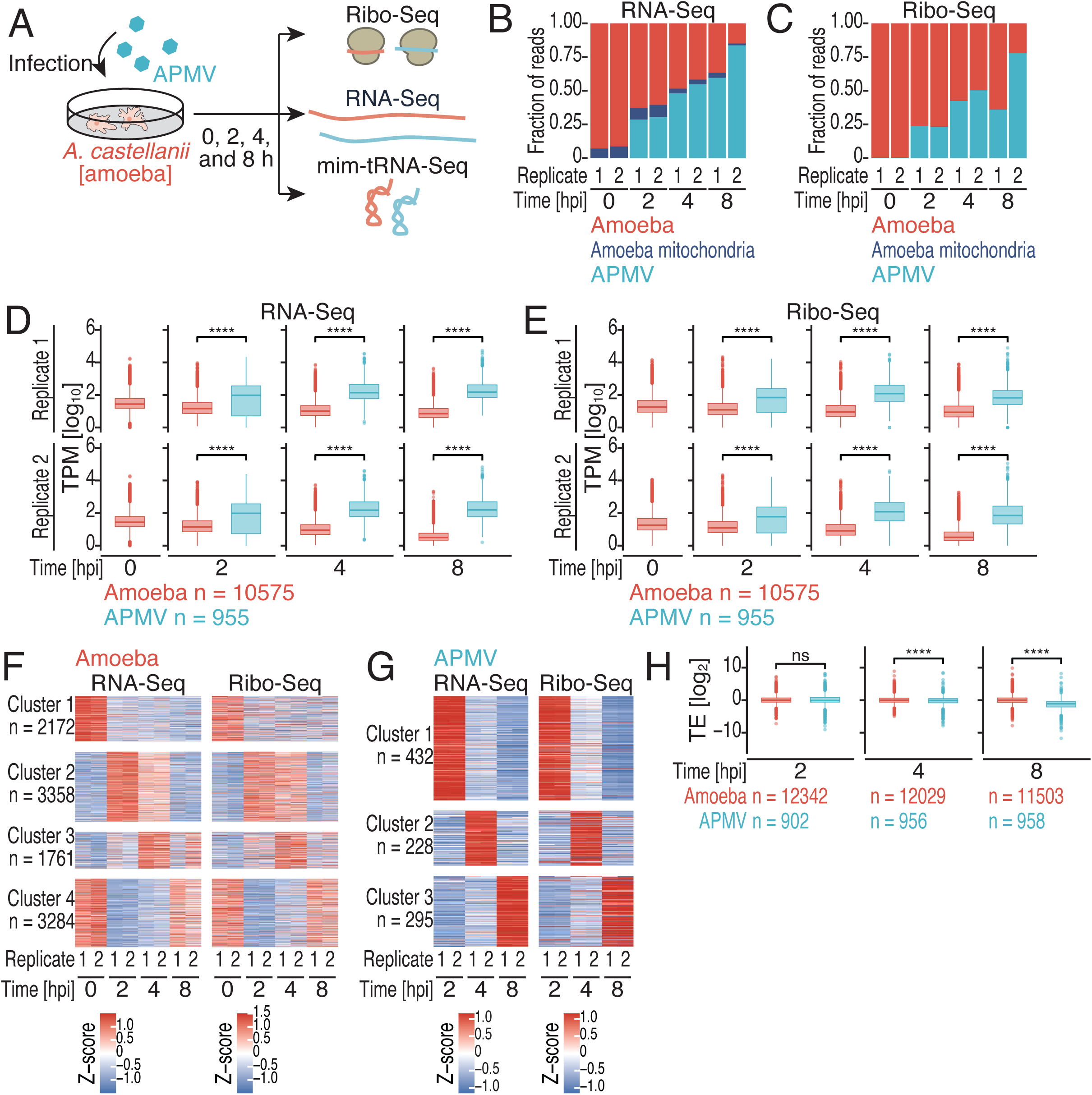
Global impacts of APMV infection on RNA abundance and translation. (A) Schematic diagram of the experimental design. Cells were infected with APMV for 0 (mock infection), 2, 4, or 8 h and used for RNA-Seq, Ribo-Seq, and mim-tRNA-Seq. (B, C) Fraction of reads that aligned to indicated genomes by RNA-Seq (B) and Ribo-Seq (C) analyses at different infection stages. hpi, hours post-infection. (D, E) Box plots of transcripts per kilobase million (TPM) for amoeba and APMV mRNAs in the RNA-Seq (D) and Ribo-Seq (E) data. (F, G) Heatmaps of the Z-score normalized TPM in the RNA-Seq and Ribo-Seq data for amoeba (F) and APMV (G) mRNAs at different infection stages. Colour scales indicate the Z-scores. (H) Box plots of translation efficiency of amoeba and APMV mRNAs at different infection stages. The median (centre line), upper/lower quartiles (box limits), 1.5× interquartile range (whiskers), and outlier (points) are shown. (D, E, H) The *p* values were calculated by the one-sided Wilcoxon rank sum test. ns, not significant; *, *p* <0.05; **, *p* <0.01; ***, *p* <0.001; ****, *p* <0.0001. See also Extended Data Fig. 1 and 2.

The RNA-Seq reads from viral mRNAs increased incrementally along with the viral infection time and were dominant at the late time points (up to approximately 62% and 85% for each replicate at 8 hpi) (Fig. 1B). Footprints from viral mRNAs also increased and occupied a large proportion of the library at 8 hpi (approximately 40% and 81% for each replicate) (Fig. 1C). We also observed a general increase of short footprints during the course of virus infection (Extended Data Fig. 2I, J).

The average expression levels of viral mRNAs were higher than those of amoeba mRNAs at all time points (even at 2 hpi) (Fig. 1D). The expression levels of individual viral mRNAs increased gradually, whereas those of host mRNAs decreased over the course of infection (Fig. 1D). Translation activity, evaluated by Ribo-Seq, followed the same trends as that obtained by RNA-Seq (Fig. 1E).

The host genes were classified into four clusters based on their mRNA levels from RNA-Seq (Fig. 1F). To characterize the functions of genes in each cluster, we selected host genes that had high variance in their expression and performed a functional enrichment analysis using Kyoto Encyclopedia of Genes and Genomes (KEGG) (Extended Data Fig. 3). Genes in cluster 1, which showed an expression shutdown by 2 hpi, were enriched in energy metabolism, amino acid metabolism, and motor proteins. Ribosomal genes had various expression patterns and were enriched in clusters 1, 3, and 4. tRNA and protein processing functions were enriched in clusters 2 and 3. Viral mRNAs were classified into three expression clusters (Fig. 1G), which largely correspond to the early, intermediate, and late genes reported previously^8^. The translation levels of host and viral mRNAs in individual clusters showed dynamics consistent with the transcription patterns, suggesting that the abundance of mRNAs is a major determinant of the level of translation (Fig. 1F, G).

To assess the density of host ribosomes on host and viral mRNAs, we calculated the translation efficiency (TE) as the ratio of the amounts of footprints to the amounts of mRNAs. At 0 and 2 hpi, the median TE values were 0.991 and 0.992 for host mRNAs, respectively, and 0.935 (2 hpi) for viral mRNAs, indicating no significant difference between the TE of host and viral mRNAs (Fig. 1H). However, viral TE dropped and was significantly lower than that of host TE at later time points; at 4 and 8 hpi, the median TE values were 0.989 and 1.010 for host mRNAs, and 0.956 and 0.468 for viral mRNAs (Fig. 1H). This result implies that host mRNAs were twice as likely to bind to ribosomes as viral mRNAs at 8 hpi, indicating the reduced accessibility to ribosomes for viral mRNAs at later time points.

### Translation of viral mRNAs is not associated with ribosome pausing

Given the deviated codon usage of APMV mRNAs, we investigated whether the rare codons on viral mRNAs hampered virus mRNA translation. Since APMV genes are biased to codons ending with A or U (hereafter AU3 codons) and host genes are biased to codons ending with G or C (hereafter GC3 codons) (Extended Data Fig. 1A), we investigated the translation elongation speed on codons from viral and host mRNAs by calculating ribosome occupancy. No significant difference was found in ribosome occupancy at the A-site between AU3 and GC3 codons for amoeba (Fig. 2A), and this balanced elongation was maintained in viral mRNAs (Fig. 2B).

**Fig. 2.**
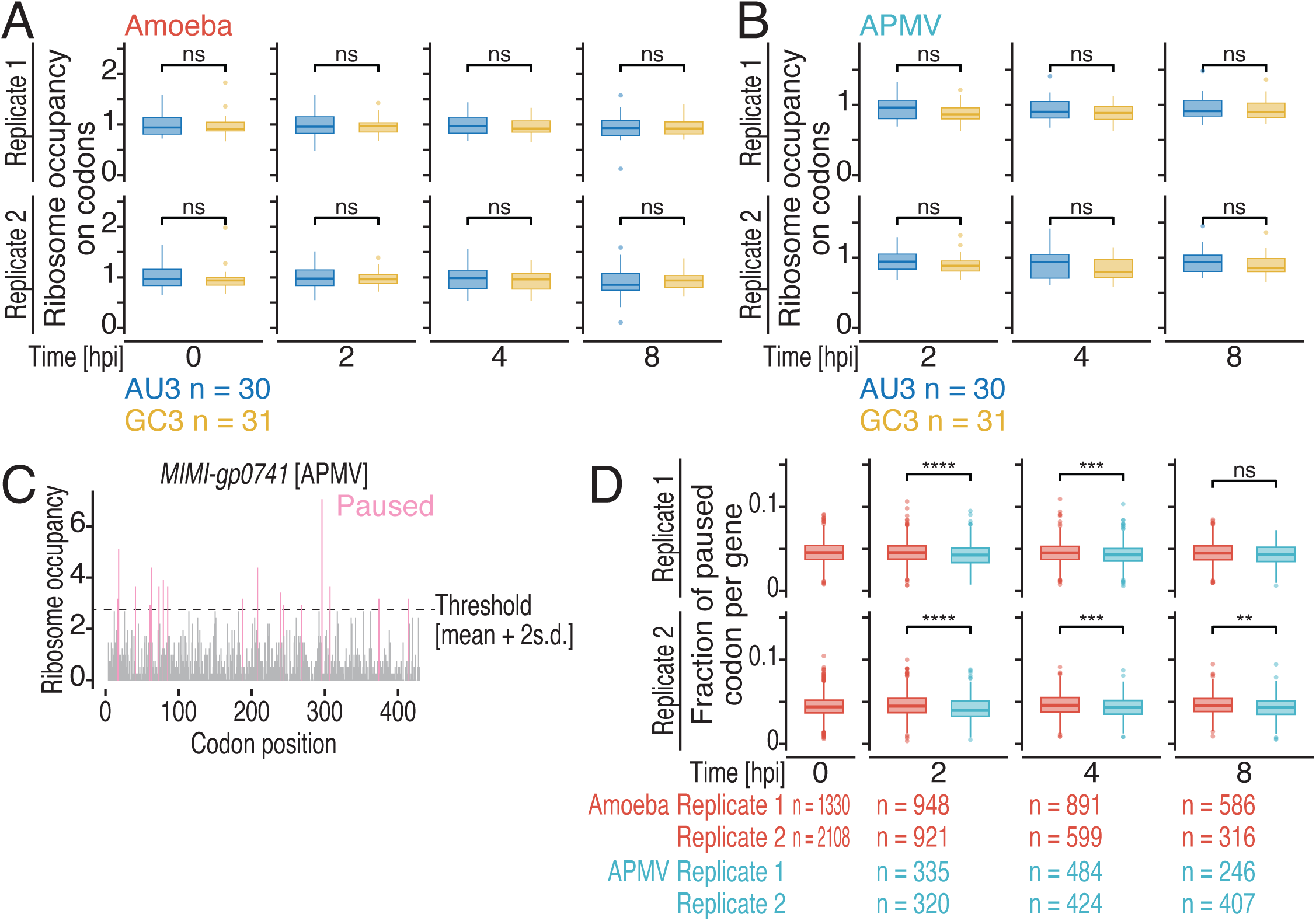
Smooth ribosome traversal on APMV mRNAs. (A, B) Box plots of ribosome occupancy on AU3 and GC3 codons in amoeba (A) and APMV (B) mRNAs at different infection stages. AU3, codons ending with A or U; GC3, codons ending with G or C; hpi, hours post-infection. (C) Distribution of ribosome footprints along the open reading frames of *MIMI-gp0741* (locus tag of an APMV gene) in the APMV genome for replicate 1 at 2 hpi. The A-site position of the ribosomes is shown. The first four codons and stop codons were not included and are not plotted. The grey horizontal line (Threshold) indicates the cutoff of pausing [mean + two standard deviations (2 s.d.)]. Pause sites are shown in pink. (D) Box plots of fraction of paused codons per gene in the amoeba and APMV genomes at different infection stages. The median (centre line), upper/lower quartiles (box limits), 1.5× interquartile range (whiskers), and outlier (points) are shown. (A, B, D) The *p* values were calculated by the one-sided Wilcoxon rank sum test. ns, not significant; *, *p* <0.05; **, *p* <0.01; ***, *p* <0.001; ****, *p* < 0.0001. See also Extended Data Fig. 2.

Next, we focused on individual codon positions to determine whether ribosomes tended to pause on either viral or host mRNAs. Codon positions with relatively high ribosome occupancy were defined as pause sites (Fig. 2C, see Methods for details). We found that viral mRNAs had a lower tendency of ribosome pausing than host mRNAs did (Fig. 2D), indicating that ribosome traversal on viral mRNAs was smooth, even with the obvious codon usage conflict (Extended Data Fig. 1).

### tRNA pool was stable during viral infection

Because APMV encodes tRNA genes [three Leu (two TTAs, one TTG), Trp (TGG), Cys (TGC), and His (CAC)]^13^, we hypothesized that alteration of the cellular tRNA pool facilitated ribosome traversal along viral mRNAs. To assess the abundance of tRNA species, we applied modification-induced misincorporation tRNA sequencing (mim-tRNA-Seq)^17^. As tested in an earlier study^17^, we also evaluated the salt concentration, pH, temperature, and reverse transcription enzymes for optimal cDNA synthesis (Extended Data Fig. 4A, B). Ultimately, we selected the most efficient condition (with the TGIRT-III enzyme at 49°C for 16 h) for downstream tRNA sequencing.

The mim-tRNA-Seq showed that the expression of tRNAs encoded in the amoeba genome remained at a high level (>87% in every library), whereas the expression of the six tRNAs encoded by APMV was only 1.8%, even at the late infection stage (8 hpi) (Fig. 3A), suggesting that the contribution of viral tRNA to the cellular tRNA pool may be limited. In addition, the codons decoded by the APMV-encoded tRNAs did not show significantly different ribosome occupancy (i.e., elongation speed) compared to other codons in viral mRNAs (Extended Data Fig. 5).

**Fig. 3.**
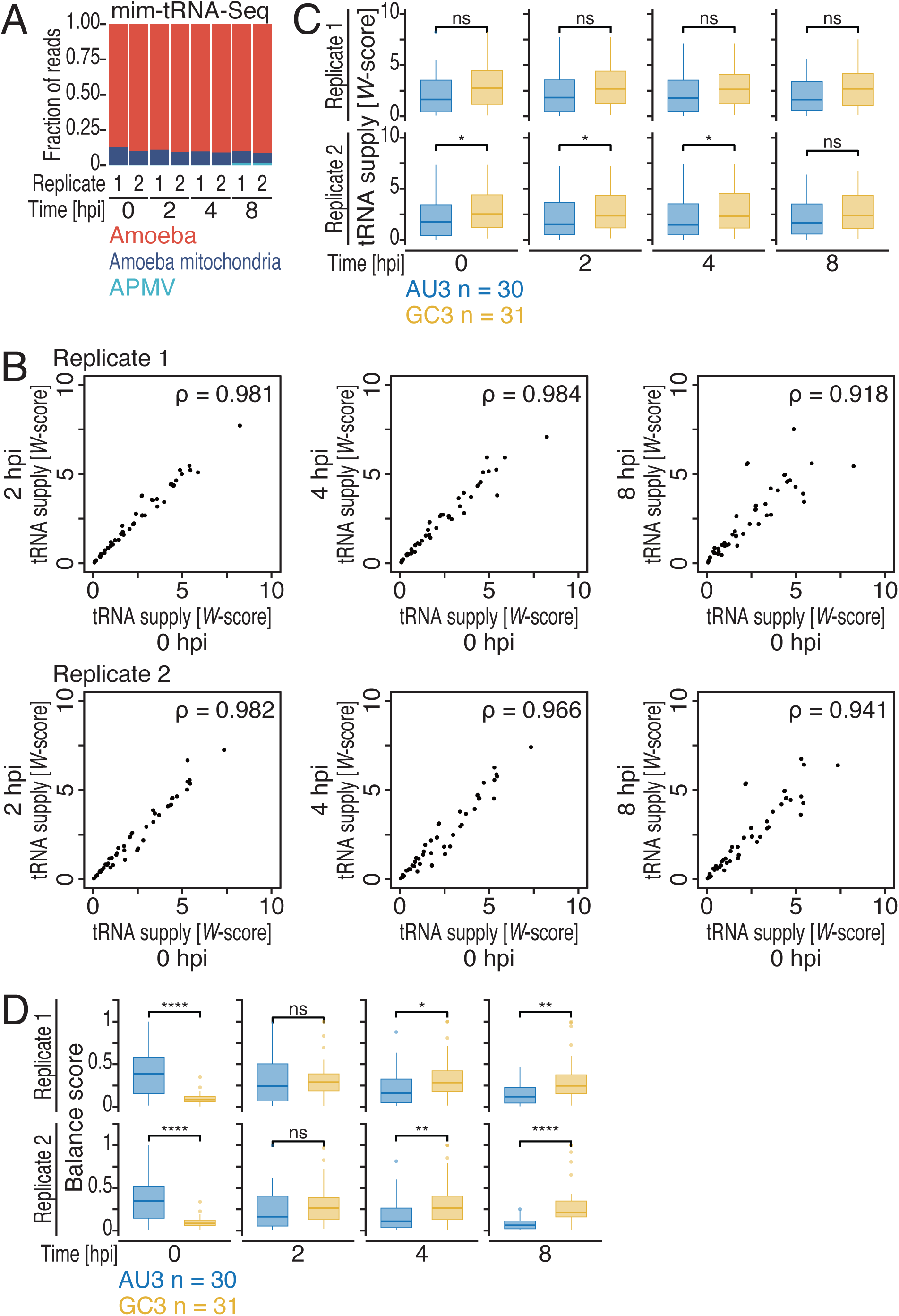
Stable tRNA pool during infection conflicts with translation elongation on APMV mRNAs. (A) Fraction of reads that aligned to the indicated genomes in the mim-tRNA-Seq analysis at different infection stages. hpi, hours post-infection. (B) Correlation of tRNA supply evaluated by *W*-score among the different infection stages. ρ, Spearman’s correlation coefficient (two-tailed). (C) Box plots of *W*-scores for AU3 and GC3 codons at different infection stages. AU3, codons ending with A or U; GC3, codons ending with G or C. (D) Box plots of balance scores for tRNA for AU3 and GC3 codons at different infection stages. The median (centre line), upper/lower quartiles (box limits), 1.5× interquartile range (whiskers), and outlier (points) are shown. (C, D) The *p* values were calculated by the one-sided Wilcoxon rank sum test. ns, not significant; *, *p* <0.05; **, *p* <0.01; ***, *p* <0.001; ****, *p* <0.0001. See also Extended Data Fig. 2I, J.

To investigate whether the virus infection altered the tRNA pool composition to favour viral mRNA translation, we calculated the proportion of each tRNA type in the tRNA pool (including amoeba nuclear and APMV tRNAs). We used *W*-score to quantify the tRNA supply level. For a given codon type, *W*-score was calculated by summing the proportions of cognate and wobble-paired tRNAs (see Methods for details)^21^. Essentially, *W*-scores were stable during the APMV infection; high correlation of the *W*-scores was observed between time points (Fig. 3B). In naïve amoeba (i.e., 0 hpi), the *W*-scores were higher for GC3 than they were for AU3 codons (Fig. 3C & Extended Data Table 1). This finding corresponds with codon usage in the amoeba genome, which is biased to GC3 codons (Extended Data Fig. 1) and probably contributed to the smooth elongation of GC3 codons (Fig. 2A) with no shortage of tRNA supply. The high tRNA supply for GC3 codons was maintained throughout the infection, indicating that viral infection did not largely change the composition of the global tRNA pool.

To quantitatively assess the imbalance of tRNA supply and demands during APMV infection, we calculated the balance score, which borrowed the idea of the normalized translation efficiency (nTE) index^22^. Briefly, we calculated the ratio between tRNA supply and codon usage considering the mRNA abundance in the cell (see Methods for details). In naïve amoeba, the AU3 codons generally had a higher balance score than the GC3 codons (Fig. 3D), indicating that tRNAs for AU3 codons are “excessive” despite their concentration being lower than that for GC3 codons. During APMV infection, the high expression of viral mRNAs led to a decrease in the balance score for AU3 codons, whereas the balance score for GC3 increased (Fig. 3D & Extended Data Table 2). Together, these findings indicate that the tRNA supply and demands inside the cell seemed unsuitable for the translation of AU3-rich viral mRNAs (Fig. 3), even though protein synthesis from viral mRNAs was rather smooth, not hampered (Fig. 2D).

### Distinct ribosome elongation environment for viral mRNAs

The conflict between tRNA supply and demand for viral mRNA translation is based on our assumption that cellular resources, such as tRNAs, are equally available for host and viral mRNAs. Thus, our results suggest that the translation environment for viral mRNAs was not the same as that for host mRNAs. Consistent with this scenario, we observed differences in tRNA accessibility in host and viral mRNAs. Given that long (29–30 nt) and short (21–22 nt) footprints represent ribosomes with or without tRNA at the A-site^20^, we defined the short footprint ratio as the ratio of short footprints to total footprints for a given codon type to represent tRNA accessibility. The short footprint ratio was higher (i.e., lower tRNA accessibility) on AU3 codons than it was on GC3 codons in amoeba mRNAs, irrespective of the viral infection stage (Fig. 4A). This finding is consistent with the lower tRNA supply for AU3 codons compared with the tRNA supply for GC3 codons (Fig. 3C). Notably, there was no such difference in the short footprint ratio for viral mRNAs (Fig. 4B). The correlation between viral codon occupancy and host codon occupancy decreased at the late infection stage (8 hpi) (Extended Data Fig. 5), indicating divergence in codon-specific elongation speed between viral and host mRNAs. These data suggest that the translation environment of viral mRNAs differed from that of amoeba mRNAs.

**Fig. 4.**
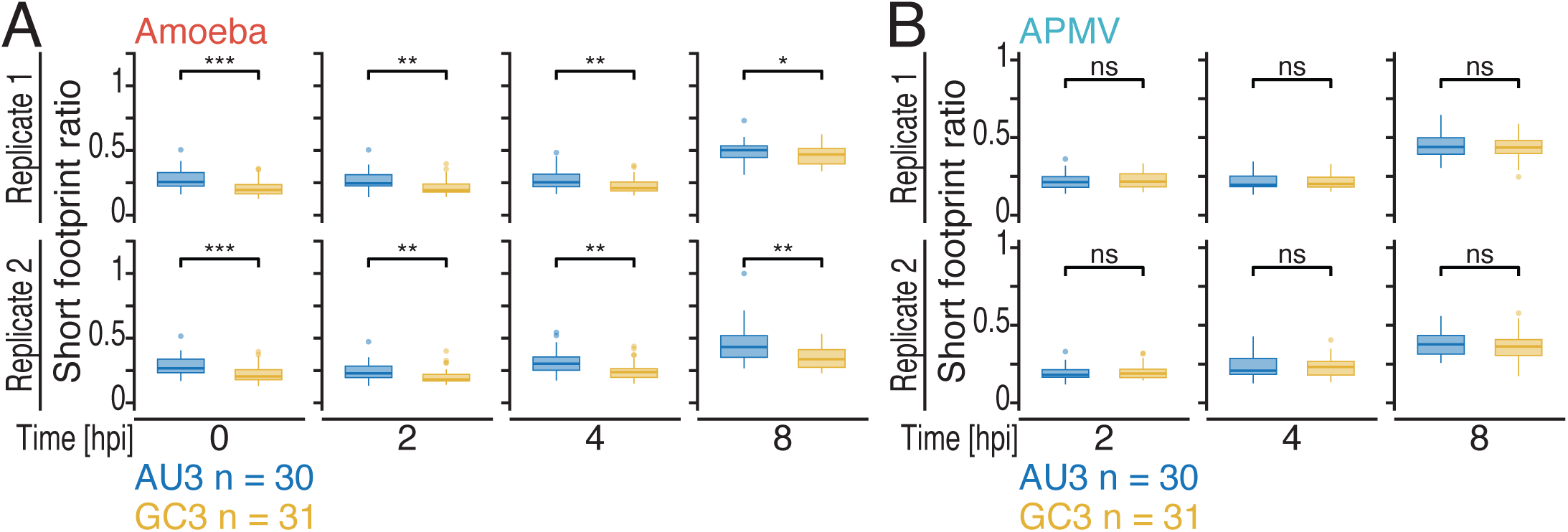
Difference in tRNA accessibility between amoeba and APMV mRNAs. (A, B) Box plots of the short footprint ratio on AU3 and GC3 codons in amoeba (A) and APMV (B) mRNAs at different infection stages. AU3, codons ending with A or U; GC3, codons ending with G or C; hpi, hours post-infection. The median (centre line), upper/lower quartiles (box limits), 1.5× interquartile range (whiskers), and outlier (points) are shown. The *p* values were calculated by the one-sided Wilcoxon rank sum test. ns, not significant; *, *p* <0.05; **, *p* <0.01; ***, *p* <0.001; ****, *p* <0.0001.

### Viral mRNAs are locally translated

As ribosomes were found to localize around the VF^15,16^, we hypothesized that viral mRNA translation may occur in a distinct subcellular location near the VF. To investigate this hypothesis, we combined 4′,6-diamidino-2-phenylindole (DAPI) staining, FISH, and FUNCAT^19^ to determine the location of the VF, viral mRNAs, amoeba ribosomes, and newly synthesized proteins. The FUNCAT signal was enriched around the periphery of the VF (Fig. 5A). Similarly, FISH for amoeba rRNA showed that a sub-population of ribosomes was also enriched in this region (Fig. 5A), which is consistent with previous tomography results^16^, suggesting that protein synthesis occurred at the VF periphery. Moreover, this local translation milieu contained the APMV-encoded *mcp* mRNAs, probed by single-molecule mRNA FISH (sm-mRNA FISH) (Fig. 5B). Thus, our data suggest that APMV created a unique environment tailored for viral mRNA translation at the periphery of the VF.

**Fig. 5.**
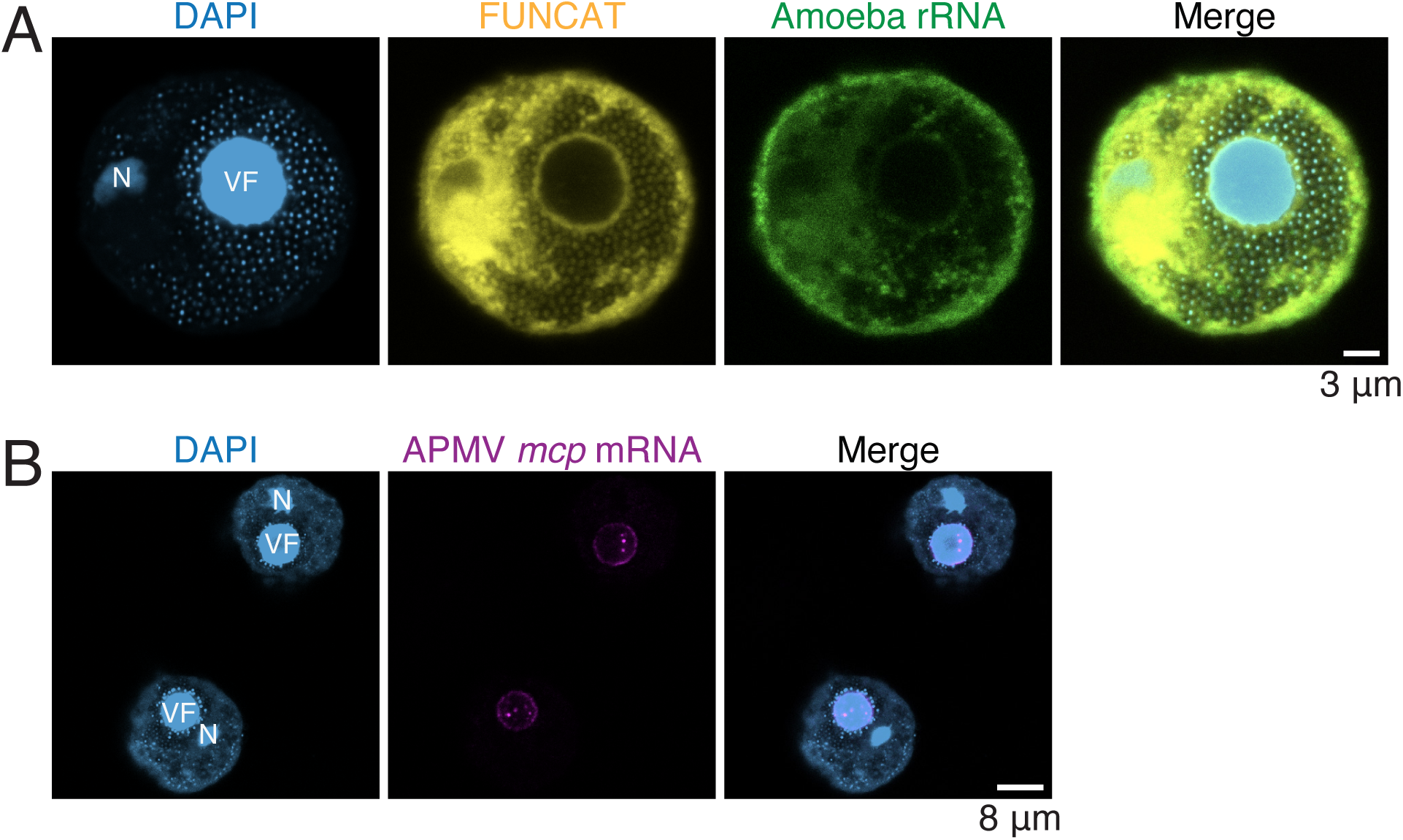
Local translation at peripheral region of the viral factory during APMV infection. (A) Microscopy images for DAPI (light blue), FUNCAT (yellow), and rRNA FISH (green) of APMV-infected amoeba cells at 12 hours post infection (hpi). Scale bar, 3 µm. (B) Microscopy images for DAPI (light blue) and single-molecule mRNA FISH for APMV *mcp* mRNA of APMV-infected amoeba cells at 18 hpi. Scale bar, 8 µm. VF, viral factory; N, amoeba nucleus.

## Discussion

A consonant pattern between codon usage and tRNA concentration was thought to be important for translation^2,3^. However, many eukaryotic viruses, including APMV, have codon usage patterns that deviate from those of their hosts^23–25^, suggesting potential problems in viral gene translation. The deep sequencing-based techniques (RNA-Seq, Ribo-Seq, and mim-tRNA-Seq) showed that APMV genes were smoothly translated without reduced accessibility to tRNAs for frequently used codons. The cell biology methods (FUNCAT and FISH) showed that viral mRNAs, host rRNA, and protein synthesis were co-localized at the periphery of VF. These findings suggest that APMV mitigates the codon usage conflict by creating a local translation milieu for its mRNAs.

Despite that APMV encodes six tRNA genes, our data indicate that APMV infection did not significantly alter the tRNA composition in the host cell (Fig. 3). This unaltered cellular tRNA pool aligns with the previous report on vaccinia virus and influenza A virus^23^, implying that a local tRNA pool could be selectively used for viral mRNAs. APMV may also achieve the smooth translation by generating a local translation environment within the cell.

The VF is a unique intracellular structure that is formed by many large DNA viruses during infection. For APMV, DNA replication and transcription occur in the VF^26,27^, whereas ribosomes and the viral translation initiation factor SUI1 have been observed outside the VF^15,16,27^. In this study, we showed that viral mRNAs surrounded the VF (Fig. 5B) and protein synthesis occurred in the same region (Fig. 5A). Other translation-related proteins and tRNAs may also be recruited to the same place, which would further increase the local concentration of translation machineries. Although viral mRNAs dominated the mRNA pool at the late stages of infection, this increase was not perfectly reflected in the footprint proportion (Fig. 1B, C), leading to a decline in TE of viral mRNAs (Fig. 1H). Similar results have been reported during infections with influenza A virus and coronavirus SARS-CoV-2^28,29^. These data suggest an overload of viral mRNAs compared with the cellular resources. Consistently, we found a general increase of A-site free ribosomes, represented by short footprints along the viral infection stages (Extended Data Fig. 2A, B and Extended Data Fig. 2I, J), probably because of the global tRNA or amino acid shortage.

Why does APMV maintain the AT-rich genome despite it conflicts with amoeba tRNA supply? Given that giant viruses enriched with translation-related genes have an AT-rich genome^30^, the preference for the AT-rich genome in APMV may be an adaptive strategy. Using AU3 codons may be advantageous in competing with amoeba mRNAs at the early stage of infection when there is an excess of tRNAs for AU3 codons (Fig. 3D). This advantage, however, is likely to cease at later infection stages with high viral mRNA load (Fig. 3D). APMV alleviates this translation-associated bottleneck by establishing a dedicated local environment to isolate a sub-population of tRNAs and other translation factors to avoid competition for resources with amoeba mRNAs. This translation-associated bottleneck may drive the continuous acquisition of translation-related genes by the AT-rich genomes of APMV-related viruses.

## Methods

### Cells and viruses

*Acanthamoeba castellanii*, strain Neff [American Type Culture Collection (ATCC), 30010] was cultured in peptone-yeast-glucose (PYG) medium at 28°C. Although a recent study proposed a reclassification of this strain^31^, it has not yet been recognized by NCBI. We used the original naming for consistency with the NCBI database. Acanthamoeba polyphaga mimivirus (APMV) was used as a prototype of mimiviruses^32^. The virus titre was determined by the 50% tissue culture infectious dose (TCID50) assay using *A. castellanii* cells.

### Lysate preparation

Lysate for Ribo-Seq, RNA-Seq, and mim-tRNA-Seq was prepared as described previously^33^. Briefly, *A. castellanii* cells were seeded on 10-cm plates at 3 × 10^6^ cells/ml supplemented with 10 ml of PYG medium. Amoeba cells were infected with APMV at a multiplicity of infection (MOI) of 10. The plates were incubated for 1 h at room temperature to allow viral absorption, then the medium was replaced with fresh PYG medium. The medium replacement timing was set to 0 hpi, and APMV-infected cells were collected at 2, 4, and 8 hpi. For mock treatment, PYG medium was used instead of viral suspension and collected at 0 hpi. Two independent replicates were prepared for each time point.

Cells were treated with 100 µg/ml cycloheximide for 1 min, then scraped down and pelleted at 300 × *g* at 4°C for 3 min, washed with 1 ml of PBS containing 100 µg of cycloheximide (Sigma– Aldrich) and 100 µg of chloramphenicol (Wako Pure Chemical Industries, Ltd.), and pelleted again at 300 × *g* at 4°C for 2 min. The cell pellets were resuspended with 600 µl lysis buffer (20 mM Tris pH 7.5, 150 mM NaCl, 5 mM MgCl_2_, 1 mM dithiothreitol, 1% Triton X-100, 100 µg/ml of cycloheximide, and 100 µg/ml chloramphenicol), treated with 5 µl of TURBO DNase (2 U/µl, Thermo Fisher Scientific) on ice for 10 min, then centrifuged at 20,000 × *g* at 4°C for 10 min. The supernatant was stored at −80°C before library construction. RNA concentration of cell lysates was measured with a Qubit RNA BR Assay Kit (Thermo Fisher Scientific).

### Preparation of Ribo-Seq library

Ribosome profiling was performed as described previously^33^. The cell lysate containing 10 µg total RNA was incubated with 2 U of RNase I (LGC Biosearch Technologies) in a 300-µl reaction mixture at 25°C for 45 min. The RNase digestion was stopped by adding 10 µl of SUPERase•In (Thermo Fisher Scientific). Ribosomes were isolated by sucrose cushion ultracentrifuged at 100,000 rpm (543,000 *× g*) for 1 h at 4°C with a TLA110 rotor and an Optima MAX-TL ultracentrifuge (Beckman Coulter). Subsequently, RNA was purified with TRIzol-LS reagent (Thermo Fisher Scientific) and a Direct-zol RNA Microprep kit (Zymo Research). RNA fragments of 17–35 nt were gel-purified after running polyacrylamide gel electrophoresis. The isolated RNA fragments were ligated to custom-made pre-adenylated linkers containing unique molecular identifiers and barcodes for library pooling using T4 RNA ligase 2, truncated KQ (New England Biolabs). Ribosomal RNA was depleted using a Ribo-Zero Gold (Human/Mouse/Rat) kit (Illumina), followed by a pull-down using RNAClean XP beads (Beckman Coulter). The rRNA-depleted samples were reverse transcribed with ProtoScript II (New England Biolabs), circularized with CircLigase II (LGC Biosearch Technologies), and PCR-amplified using Phusion polymerase (New England Biolabs). The libraries were sequenced on an Illumina HiSeqX system (Illumina) with the 150 nt paired-end read option.

### Preparation of RNA-Seq library

Total RNA was purified from the cell lysate using TRIzol-LS and a Direct-zol RNA Microprep kit (Zymo Research). RNA-Seq libraries were prepared with TruSeq Stranded Total RNA Library Prep Gold (Illumina). The library was sequenced on a HiSeqX system (Illumina) with the 150 nt paired-end reads option.

### Preparation of mim-tRNA-Seq library

The tRNA fraction was collected from the lysate using a mirVana miRNA Isolation Kit (Thermo Fisher Scientific) and deacylated by incubating at 37°C for 45 min in 100 mM Tris-HCl pH 9.0. The library preparation scheme was adapted from the method for Ribo-Seq^33^, using 80 ng of tRNAs. Then, 2 µl of 1.25 µM reverse transcription primer (NI-802)^33^ was hybridized to linker-ligated tRNA dissolved in 10 µl of RNase-free water by denaturing at 82°C for 2 min and cooling at 25°C for 5 min in a thermocycler. Reverse transcription was conducted in 21.6 mM Tris-HCl pH 7.5, 5.4 mM MgCl2, and 486 mM KCl containing 5.4 mM dithiothreitol, 0.54 U/µl SUPERase•In (Thermo Fisher Scientific), and 540 nM TGIRT-III (InGex) in an 18.5-µl reaction mixture and pre-incubated at 42°C for 10 min in a thermocycler. Next, 2.5 µl of 10 mM dNTP was added to the tube to make a 20-µl reaction mixture and incubated at 49°C for 16 h in a thermocycler. Subsequent circularization and PCR amplification were conducted as in the Ribo-Seq^33^. The library was sequenced on a HiSeqX system (Illumina) with the 150 nt paired-end reads option.

### Construction of non-coding RNA database

We constructed a non-coding RNA (ncRNA) database by collecting annotated sequences from databases and *de novo* prediction. The annotated ncRNA sequences of *A. castellanii* were downloaded from the RNAcentral database (17 August 2023) and searched by queries *Acanthamoeba castellanii*, “TAXONOMY 5755”, and “TAXONOMY 1257118” (RNAcentral Consortium, https://rnacentral.org)^34^. The “TAXONOMY 5755” search identified 241 ncRNA records; 205 rRNAs, 32 sncRNAs, 3 RNase P RNAs, 1 RNase MRP RNA, and 1 group I intron. The “TAXONOMY 1257118” search identified 205 ncRNA records; 44 rRNAs, 149 sncRNAs, 1 RNase P RNA, 1 RNase MRP RNA, 4 SRP RNAs, and 2 vault RNAs.

For *de novo* prediction, the nuclear genome sequence of *A. castellanii* was downloaded from the latest chromosome-scale genome assembly^14^. The mitochondrial (NC_001637.1) and APMV (NC_014649.1) genome sequences were downloaded from the RefSeq database and used for *de novo* ncRNA prediction. Ribosomal RNA genes were predicted using RNAmmer v.1.2^35^, and tRNA genes were predicted using tRNAscan-SE v.2.0.12 (parameters: -HQ for amoeba, -G -HQ for APMV, -mt -Q for mitochondria)^36^.

### Read mapping

For the Ribo-Seq and RNA-Seq reads, data were processed as described previously^37^. Briefly, adapter trimming was performed with fastp v.0.21.0^38^. Reads were mapped to the ncRNA sequences using STAR v.2.7.0a^39^ to remove reads that originated from rRNA genes and other ncRNA genes *in silico*. The remaining reads were mapped to the merged genomes of the virus, amoeba nucleus and mitochondria, using STAR v.2.7.0a. BAM file indexing and read extraction were performed using SAMtools v.1.10^40^. Offsets for footprints of different lengths were determined by checking the enrichment of footprints around the start and stop codons of genes (Extended Data Table 3).

The mim-tRNA-Seq, reads were quality-controlled using fastqc v.0.12.1 (http://www.bioinformatics.babraham.ac.uk/projects/fastqc/) and regions after 97 nt (quality) were trimmed using fastx-trimmer v.0.0.14 (http://hannonlab.cshl.edu/fastx_toolkit/). The trimmed reads were mapped to the ncRNA sequences (except for tRNAs) using Bowtie2 to remove other ncRNAs^41^. The unmapped reads were extracted and mapped to the tRNA sequences using Bowtie2.

### Gene filtering, quantification, and normalization

To quantify gene expression levels, reads per kilobase per million mapped reads (RPKM) values were calculated for protein-coding genes in the amoeba and APMV genomes based on the total number of reads that uniquely aligned to the coding regions of amoeba or virus, respectively.

Footprints that mapped to the first five codons (including the start codon) or the last five codons (including the stop codon) were not counted. Open reading frames with total RNA and footprint RPKM values <10 in any replicate were discarded. After filtering, 10,575 and 955 protein-coding genes from the amoeba nuclear genome and APMV genome, respectively, remained for subsequent analyses.

Transcripts per kilobase million (TPM) values were used to compare host and viral gene expression levels. To analyse the gene expression patterns in the host or virus, TPM values were recalculated separately for the host and virus genes. TE was calculated using DEseq2^42^ as a mean fold-change between footprint and read counts considering replicates.

### Gene clustering and enrichment analysis

After filtering lowly expressed genes, Z-score normalization was performed on each gene in one replicate. The optimal number of K-means clusters was determined based on TPM values using the R package factoextra v.1.0.7 (https://rpkgs.datanovia.com/factoextra/index.html). KEGG pathway and KEGG ortholog groups were assigned using the eggNOG-mapper webserver v.2.1.1.2^43^ (http://eggnog-mapper.embl.de/) with the following settings: e-value <1E−5, bit score ≥60, identity ≥60%, query coverage ≥20%, and subject coverage ≥20%. The KEGG Pathway database for amoeba was built with the assigned KO (KEGG Orthology) terms using a homemade R code. Enrichment analyses were performed for the top 2000 genes with the highest standard deviation using clusterProfiler v.4.8.3^44^.

### Quantification of tRNA expression

For tRNA read mapping, reads that met the following criteria were retained: (1) length ≥20 bp; (2) uniquely mapped to tRNA genes that encode the same anticodons; (3) anticodon of mapped tRNA was not “NNN”, “TTA” (*i.e.*, tRNA-suppressor), or “TCA” (*i.e.*, tRNA for selenocysteine); and (4) reads mapped to the same genome. These criteria ensured that reads with unique mapping and reads that mapped to multiple genes but encoded the same type of anticodon were kept for quantification.

The composition of the cellular tRNA pool was calculated as the proportion of tRNA species in each sample. Because tRNA and a codon can pair by wobble pairing, we used absolute adaptiveness (*W*) to measure the tRNA supply to each type of codon^21^. The *W*-score of a codon was computed as the weighted sum of selection constraint (𝑠_𝑖𝑗_) and tRNA ratio (𝑡𝑅𝑁𝐴_𝑗_) considering both Crick’s rules and wobble pairing. The selective constraint (𝑠_𝑖𝑗_) represents the difficulty of the pairing between codon 𝑖 and anticodon 𝑗.

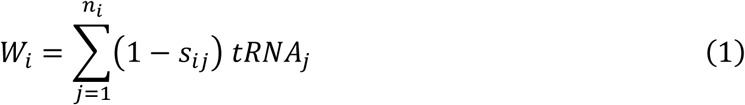

Briefly, to infer 𝑠_𝑖𝑗_, we performed Nelder-Mead optimization on the Spearman’s correlation between the expression level and computed tRNA adaptation index (*tAI*) of corresponding genes using the Python package scipy^45^. The *tAI* is a measure of the tRNA usage by coding sequences^21^. For gene 𝑔, 𝑡𝐴𝐼_𝑔_ was computed as the geometric mean of the relative adaptiveness 𝑤_𝑖_ of its codons. If 𝑡𝐴𝐼_𝑔_ = 1, it indicates gene 𝑔 is well supported by the tRNA pool, whereas 𝑡𝐴𝐼_𝑔_ close to 0 indicates that the gene is poorly supported.

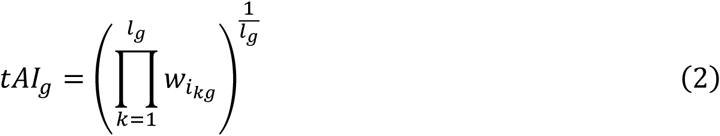

Here, 𝑖_𝑘𝑔_ is the codon at the 𝑘th position in gene 𝑔 and 𝑙_𝑔_ is the length of the gene in codons (excluding the stop codon). Consequently, 𝑡𝐴𝐼_𝑔_ estimates the amount of adaptation of gene 𝑔 to its genomic tRNA pool.

In Eq. (2), the relative adaptiveness 𝑤_𝑖_ is the normalized absolutive adaptiveness (*W*-score). The relative adaptiveness 𝑤_𝑖_ was calculated as the ratio between the 𝑊_𝑖_ for each codon and the maximum 𝑊. Thus, 𝑤_𝑖_ = 1 indicates that codon 𝑖 is the most adapted codon to the tRNA pool, and 𝑤_𝑖_ close to 0 indicates that the codon is very poorly adapted.

Originally, the maximum 𝑊 was set as the highest 𝑊 among all codons. To avoid making this value biased to being much higher than other values, we used the geometric mean of the three highest 𝑊 values (𝑊_𝑁_) as the maximum *W* value for computation. 𝑊_𝑁_was used to calculate the relative adaptiveness of each codon.

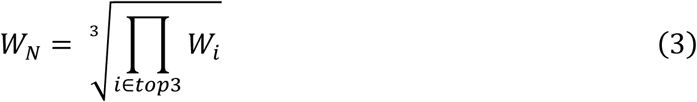

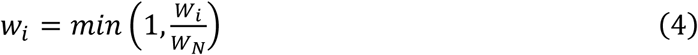

After calculating *tAI* for each gene, we performed parameter optimization for 𝑠_𝑖𝑗_ using the top 2000, 3000, 4000, and 5000 amoeba genes based on the ranking of the footprint TPM separately and initialized the parameter set from [0,0,0,0,0], [0.25, 0.25, 0.25, 0.25, 0.25], [0.5, 0.5, 0.5, 0.5, 0.5], [0.75, 0.75, 0.75, 0.75, 0.75], [1,1,1,1,1]. The 𝑠_𝑖𝑗_ set was [1, 0, 1, 0.0065956, 0.90832489] (𝜌 ≈ 0.450), which represents the wobble pairing between A:I, C:I, G:U, U:G, and A:G (for the codon–ATA:tRNA–GAT pair), respectively.

### Codon usage and balance score

For codon usage, 𝑈_𝑖_, the total number of occurrences of codon type 𝑖 was computed by summing up the occurrences of codon 𝑖 in gene 𝑔, denoted as 𝑐_𝑖𝑔_, weighted by the mRNA abundance, ^𝑇𝑃𝑀^𝑟𝑛𝑎,𝑔.

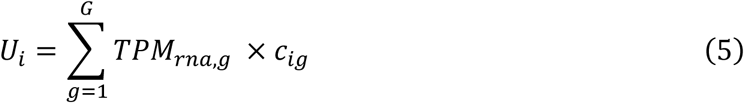

We defined codon usage 𝑐𝑢_𝑖_ as a relative estimate of how often each codon is used. We computed 𝑐𝑢_𝑖_ by rescaling 𝑈_𝑖_ to have a maximum value of 1. 𝑈_𝑚𝑎𝑥_ in Eq. (7) is the geometric mean of 𝑈_𝑖_ of the top three codons.

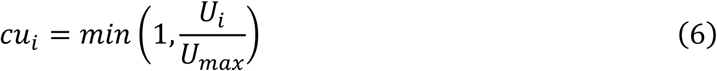

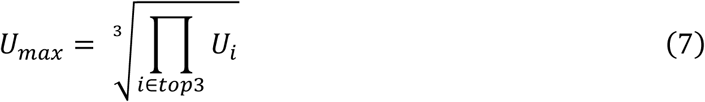

The balance score is defined as the ratio of tRNA availability, the relative adaptiveness of each codon (𝑤_𝑖_), and codon usage (𝑐𝑢_𝑖_), then linearly rescaled to have a maximum value of 1^22^.

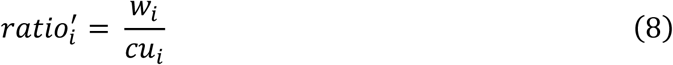

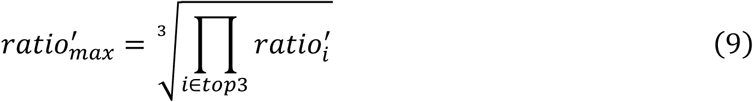

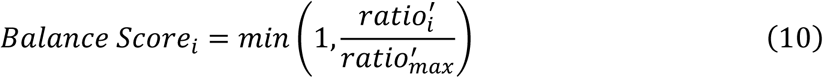

A codon with 𝑟𝑎𝑡𝑖𝑜′ >1 was defined as “excessive tRNAs” indicating that the tRNA supply was higher than the codon usage, and 𝑟𝑎𝑡𝑖𝑜′ <1 was defined as “moderate tRNAs” indicating the tRNA supply was lower than the codon usage. The balance score is derived from this score by rescaling the range from zero to one.

### Codon occupancy

The relative ribosome density at the 𝑘th codon of gene 𝑔 (𝑒_𝑘𝑔_) was defined as the number of ribosome footprints at position 𝑘 (𝑅𝑃𝐹_𝑘𝑔_) to the average number of ribosome footprints per codon of gene 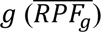.

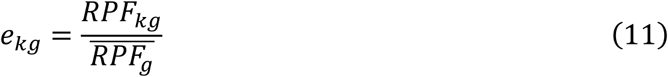

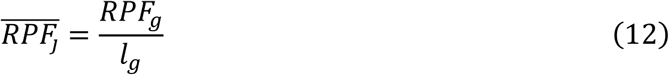

The ribosome occupancy of a given type of codon 𝑖 was the mean value of all positions where the corresponding codon is 𝑖. Only genes with 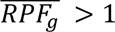 were included in this analysis. The first four codons in each gene were excluded when calculating codon occupancy^46^. Stop codons were also not included in the analysis.

### Short footprint ratio

The short footprint ratio of codon type 𝑖 was defined as the ratio between the number of short footprints (𝑅𝑃𝐹_𝑆_) mapped to codon 𝑘 and the total number of footprints (𝑅𝑃𝐹_𝑇_) mapped to codon type 𝑖. Genes that were used in the codon occupancy analysis were used in this study. The start and stop codons were excluded from the calculation.

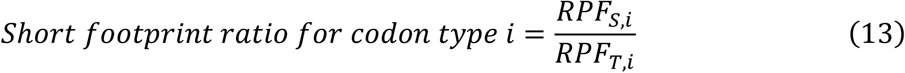

### Identification of putative pausing site

For each gene, we calculated the mean and standard deviation of the relative ribosome density as 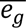 and 𝑠𝑑_𝑔_. We considered positions as putative pausing sites when 𝑒_𝑘,𝑔_ exceeds the mean 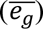 plus two standard deviations (𝑠𝑑_𝑔_) of the gene^47^. The number of putative paused sites was summed based on their codon type and the original gene. These numbers were normalized by the total occurrences of codon types and gene length excluding the first four codons and stop codon.

### Statistical test

All statistical tests were done in R v.4.3.1 (R Core Team, 2023). Significance annotation in boxplots was done using R package ggpubr v.0.6.0 (https://rpkgs.datanovia.com/ggpubr/).

### FUNCAT, single-molecule mRNA FISH, rRNA FISH, and DAPI staining

*A. castellanii*, strain Neff (ATCC 30010) cells were seeded in PYG medium at 10^5^ cells/ml in a 24-well plate (Thermo Fisher Scientific) with a final volume of 1 ml per well. The cells were incubated to attach to the dish bottom at 25°C for at least 30 min. Subsequently, the cells were infected with APMV at MOI of 1. The plates were centrifuged for 30 min at 1,000 × *g* at room temperature to synchronize the infection. Then, the medium was exchanged for fresh PYG medium to minimize the possibility of re-infections.

For each well, the 1 ml cell solution was collected and transferred into a 1.5-ml tube (Eppendorf) and centrifuged at 5,000 × *g* for 5 min at room temperature. After removing 80% of the supernatant, the remaining volume was resuspended using a vortex. Then, 50 µl of this concentrated cell suspension was added to each well of a microscopy slide (Marienfeld). The cells were incubated for 1 h to attach to the microscopy slide. The supernatant was removed, and the cells were fixed using 4% paraformaldehyde for 10 min at room temperature. The cells were washed using Milli-Q water and permeabilized with 70% ethanol for 2 h at 4°C.

FUNCAT, single-molecule mRNA fluorescence in situ hybridization (sm-mRNA FISH), and rRNA FISH were performed as described previously^48^. FUNCAT was developed from BONCAT (bio-orthogonal noncanonical amino acid tagging)^19,49–52^. Briefly, homopropargylglycine was added at a concentration of 50 µM at least 30 min before sampling. To identify newly synthesized proteins, an azide-bearing affinity tag was covalently attached by “click chemistry”. For this, 221 µl Page’s amoeba saline buffer (PAS: ATCC medium 1323) was mixed with 12.5 µl of 100 mM sodium ascorbate and 12.5 µl 100 mM aminoguanidine hydrochloride. Separately, 1.25 µl of 20 µM CuSO_4_, 1.25 µl of 100 µM Tris[(1-hydroxypropyl-1H-1,2,3-triazol-4-yl)methyl]amine (THPTA), and 0.3 µl of 5 mM of Cy3-azide dye were mixed and left to react in the dark for 3 min at room temperature. The two solutions were mixed carefully and 10 µl was added to each well of microscope slides. The slides were incubated in the dark for 30 min at room temperature, and then washed with PAS.

For the sm-mRNA FISH, 35 20-nt probes were designed to target the *mcp* gene of APMV^48^ using the Stellaris RNA FISH Probe Designer of LGC Biosearch Technologies (https://www.biosearchtech.com/support/tools/design-software/stellaris-probe-designer). To each well on the microscopy slides, 10 µl of hybridization buffer (2X SCC, 100 mg/mL Dextran sulfate, 2 mM Ribonucleoside-vanadyl complex, 0.2 mg/mL BSA, 1 mg/mL E. coli tRNA and 20% formamide) containing the probe mix (with each probe labelled with the fluorophore Cy5) was added at a working probe solution of 437.5 nM. The slides were placed in a moist chamber and incubated at 37°C for 20 h. The hybridization buffer was washed away with a wash buffer (2X SCC in nuclease-free water) at 39°C for 10 min. Then, the slides were dipped in ice-cold water and air-dried.

For the rRNA FISH, 10 µl of hybridization buffer (0.9 M NaCl, 20 mM Tris-HCl pH 8.0, 0.01% SDS, and 25% formamide) and 1 µl of Euk516 probe (5′-GGAGGGCAAGTCTGGT-3′; labelled with the fluorophore Fluos) was added to each well of the microscopy slides. Hybridization was performed in moist chambers at 46°C for 1.5 h. Subsequently, slides were washed with a wash buffer (20 mM Tris-HCl pH 8.0, 5 mM EDTA, and 0.149 M NaCl) and kept in the wash buffer in a 50 mL tube at 48°C for 10 min. Then, the slides were dipped in ice-cold Milli-Q water and air-dried.

After the sm-mRNA FISH or rRNA FISH, DAPI (1 µg/ml) was added to the wells on the slides, incubated for 3 min, and washed off with nuclease-free water and air-dried. The slides were observed using a confocal laser-scanning microscope (Leica SP8).

## Data availability

The RNA-Seq, Ribo-Seq, and mim-tRNA-Seq results obtained in this study have been deposited in the National Centre for Biotechnology Information (NCBI) database under following accession numbers.

RNA-Seq: GSE276076 (https://www.ncbi.nlm.nih.gov/geo/query/acc.cgi?acc=GSE276076)

Ribo-Seq: GSE276078 (https://www.ncbi.nlm.nih.gov/geo/query/acc.cgi?acc=GSE276078)

tRNA-Seq: GSE276080 (https://www.ncbi.nlm.nih.gov/geo/query/acc.cgi?acc=GSE276080)

## Code availability

The key codes for the analyses of RNA-Seq, Ribo-Seq, and mim-tRNA-Seq data have been deposited in Zenodo (https://doi.org/10.5281/zenodo.13748128).

## Acknowledgements

Computation time was provided by the Supercomputer System, Institute for Chemical Research, Kyoto University, and the HOKUSAI SailingShip supercomputer facility at RIKEN. This study was supported by the Japan Society for the Promotion of Science (JSPS) [JP23KJ1258 (to R.Z.), JP18H02279 (to H.O.), JP22H00384 (to H.O.), and JP24H02307 (to S.I.)], The Japan Science Society [Sasakawa Scientific Research Grant 2022-4103 (to. R.Z.)], and the Japan Science and Technology Agency (JST) [JPMJFS2123 (to R.Z.)]. A.W. acknowledges funding from the European Union’s Horizon 2020 research and innovation programme under the Marie Sklodowska-Curie grant agreement No. 891572 and the European Union (ERC, CHIMERA, 101039843). Views and opinions expressed are however those of the author(s) only and do not necessarily reflect those of the European Union or the European Research Council Executive Agency. Neither the European Union nor the granting authority can be held responsible for them. APMV was provided by Dr. Bernard La Scola, Aix-Marseille Université, Marseille, France. R.Z. is a recipient of a fellowship from JSPS (DC2). We thank Margaret Biswas, PhD, from Edanz (https://jp.edanz.com/ac) for editing a draft of this manuscript.

## Author Contributions

Conceptualization, R.Z., L.M., H.H., A.W., S.I., and H.O.;

Methodology, M.M., A.W., Y.S., and S.I.;

Formal analysis, R.Z. and L.M.;

Investigation, R.Z., L.M., H.H., and M.M.;

Writing – Original Draft, R.Z., L.M., S.I., H.O.;

Writing – Review & Editing, R.Z., L.M., H.H., Y.S., M.M., A.W., S.I., and H.O.;

Visualization, R.Z. and L.M.;

Supervision, H.H., Y.S., A.W., S.I., and H.O.;

Project administration, S.I. and H.O.;

Funding Acquisition, R.Z., A.W., S.I., and H.O.

---

**Extended Data Fig. 1.**
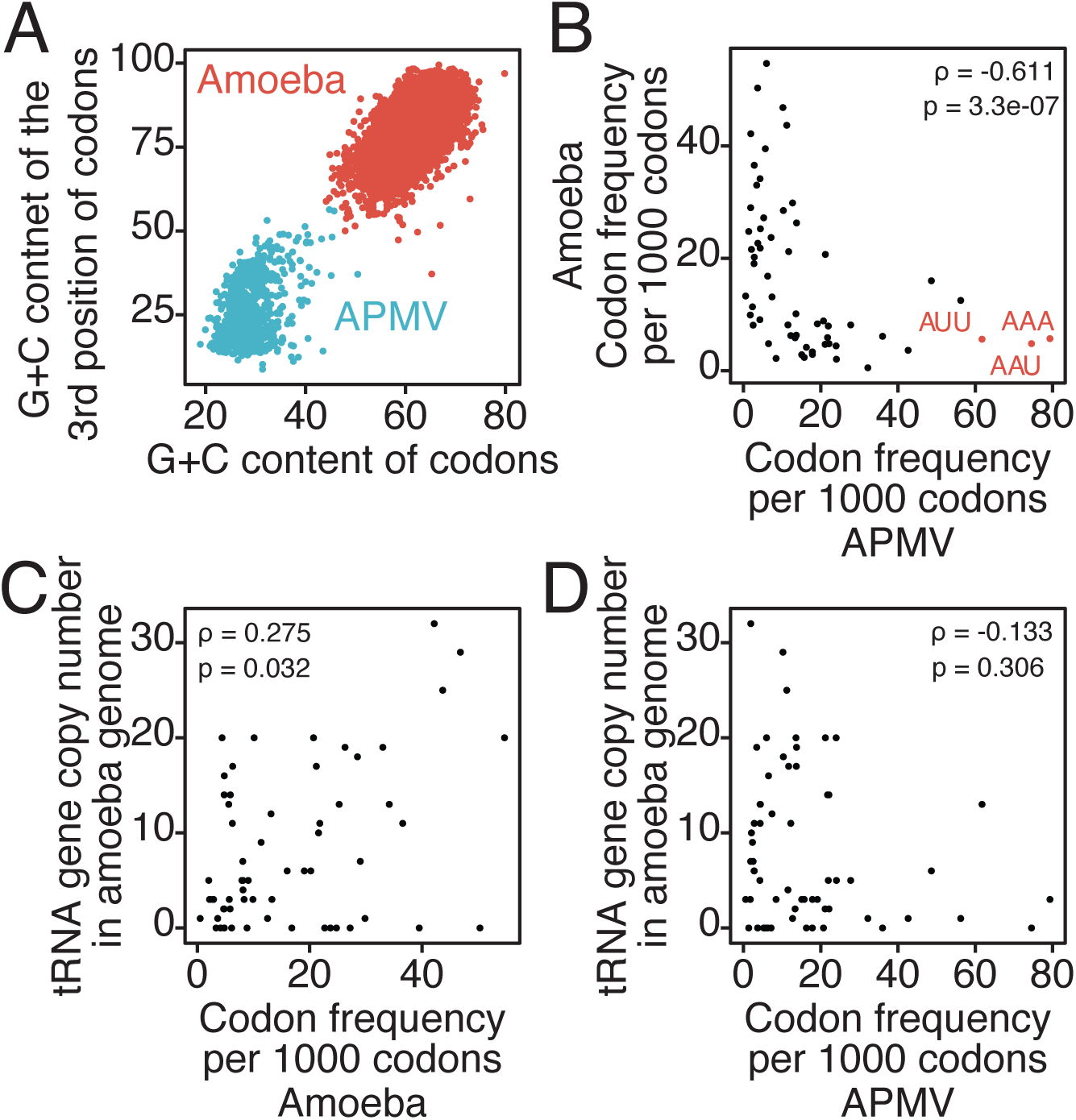
Codon usage difference between amoeba and APMV genomes (related to Fig. 1). (A) Correlation between G+C content of codons and G+C content of the 3^rd^ position of codons in amoeba (red) and APMV (light blue) mRNAs. (B) Correlation between codon frequency per 1000 codons in amoeba and APMV mRNAs. ρ, Spearman’s correlation coefficient (two-tailed); p, *p* value. (C) Correlation between codon frequency per 1000 codons of amoeba mRNAs and corresponding tRNA copy number in the amoeba genome. (D) Correlation between codon frequency per 1000 codons of APMV mRNAs and corresponding tRNA copy number in amoeba genome.

**Extended Data Fig. 2.**
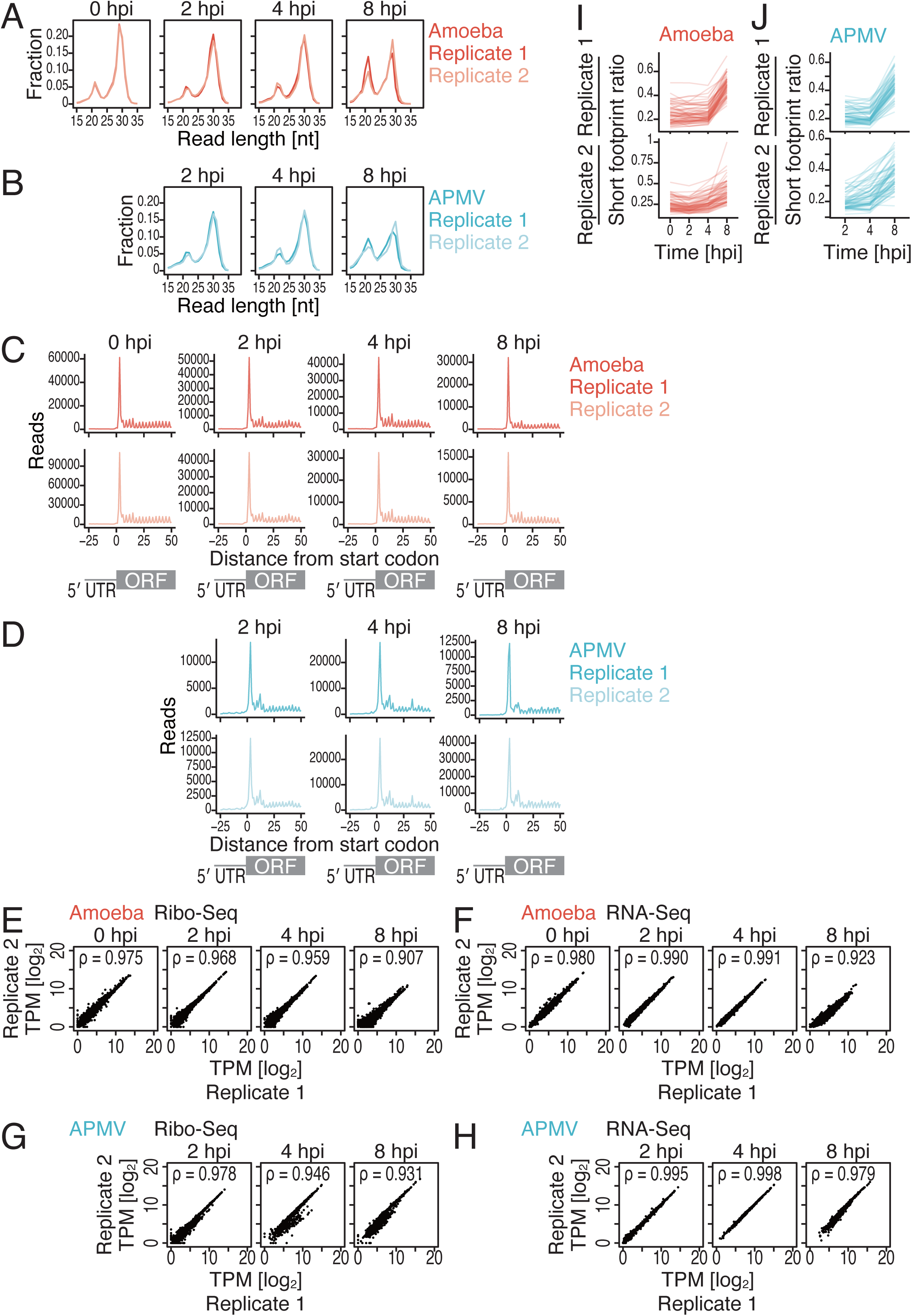
Characterization of Ribo-Seq data (related to Fig.s 1 and 2). (A, B) Distribution of ribosome footprint length at different infection stages. Reads originating from amoeba and APMV mRNAs were analysed separately. hpi, hours post-infection. (C, D) Metagene plots of the relative footprint distribution around start codons at different infection stages. Reads originating from amoeba and APMV mRNAs were analysed separately. (E–H) Scatter plots of the transcripts per kilobase million (TPM) values from two independent biological replicates for ribosomal footprints (E, G) and mRNAs (F, H) at different infection stages. (I, J) Short footprint accumulation during viral infection. Short footprint ratios on each codon type in amoeba (I) and APMV (J) mRNAs at different infection stages.

**Extended Data Fig. 3.**
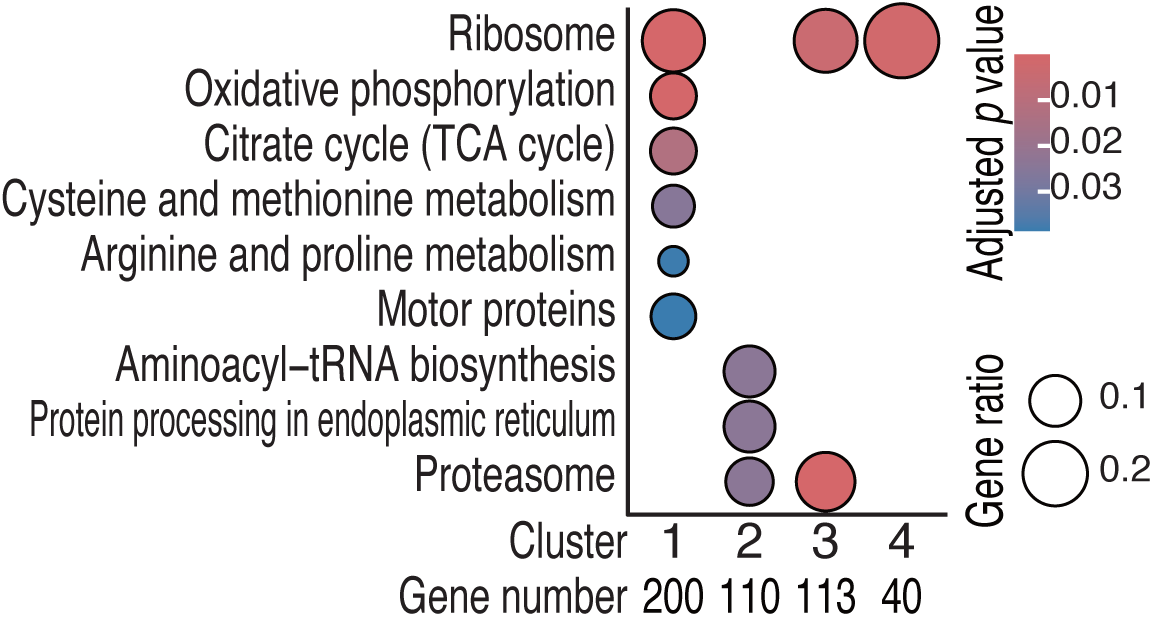
Enriched KEGG pathways in each host gene cluster. The colour scale for significance and size scale for the gene ratio in each category are shown. The *p* values were calculated by the hypergeometric distribution implemented in the enrichGO function in clusterProfiler. The Benjamini–Hochberg method was used to control the false discovery rate. The cutoffs for *p* value and *q*-value were both 0.05.

**Extended Data Fig. 4.**
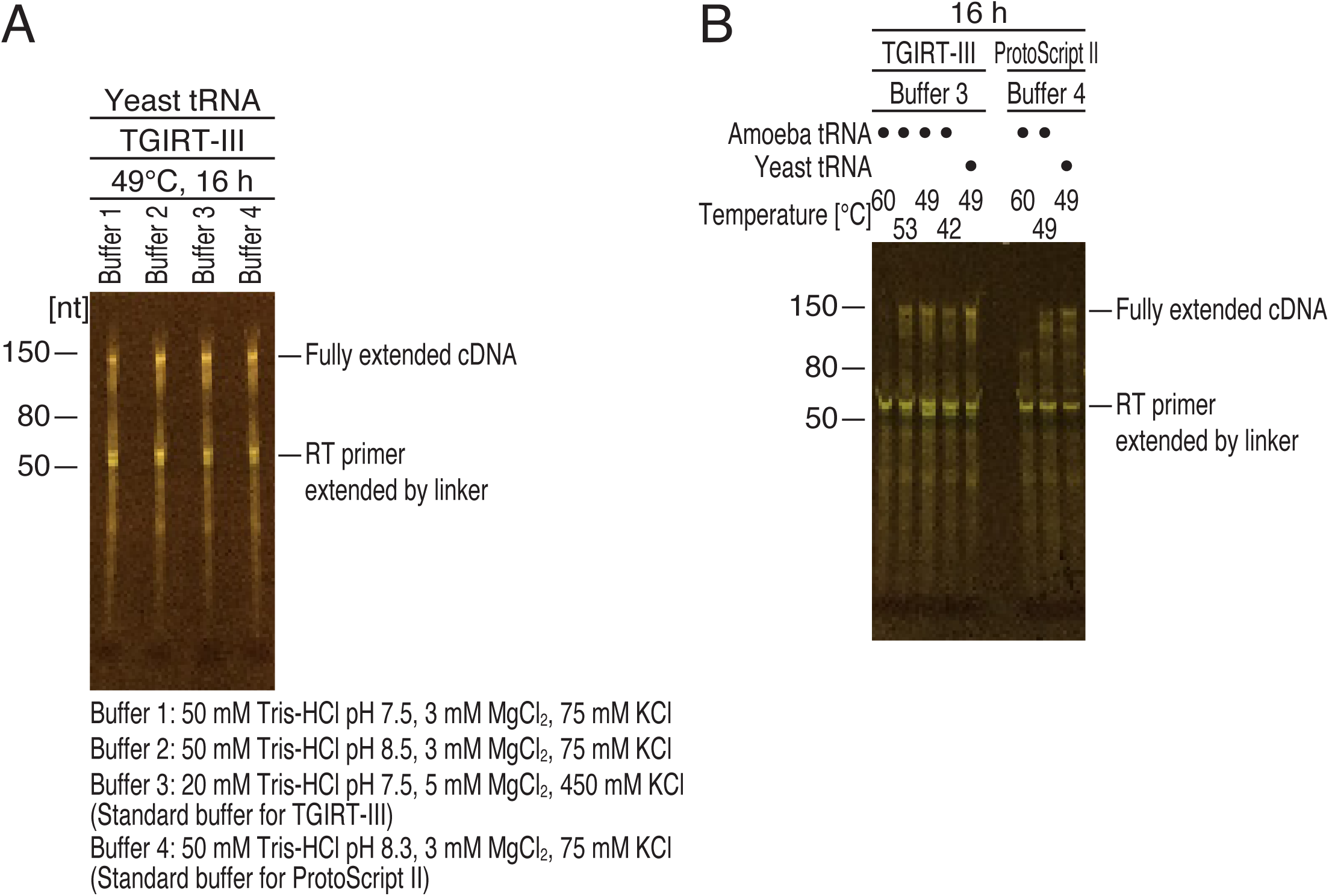
Optimization of library preparation for mim-tRNA-Seq (related to Fig. 3). (A, B) Optimization of the reverse transcription condition was conducted using titrating buffers, the reverse transcriptases TGIRT-III and ProtoScript II, and temperatures. Linker-ligated tRNAs (from yeasts or amoeba) at the 3′ end were used for the experiments.

**Extended Data Fig. 5.**
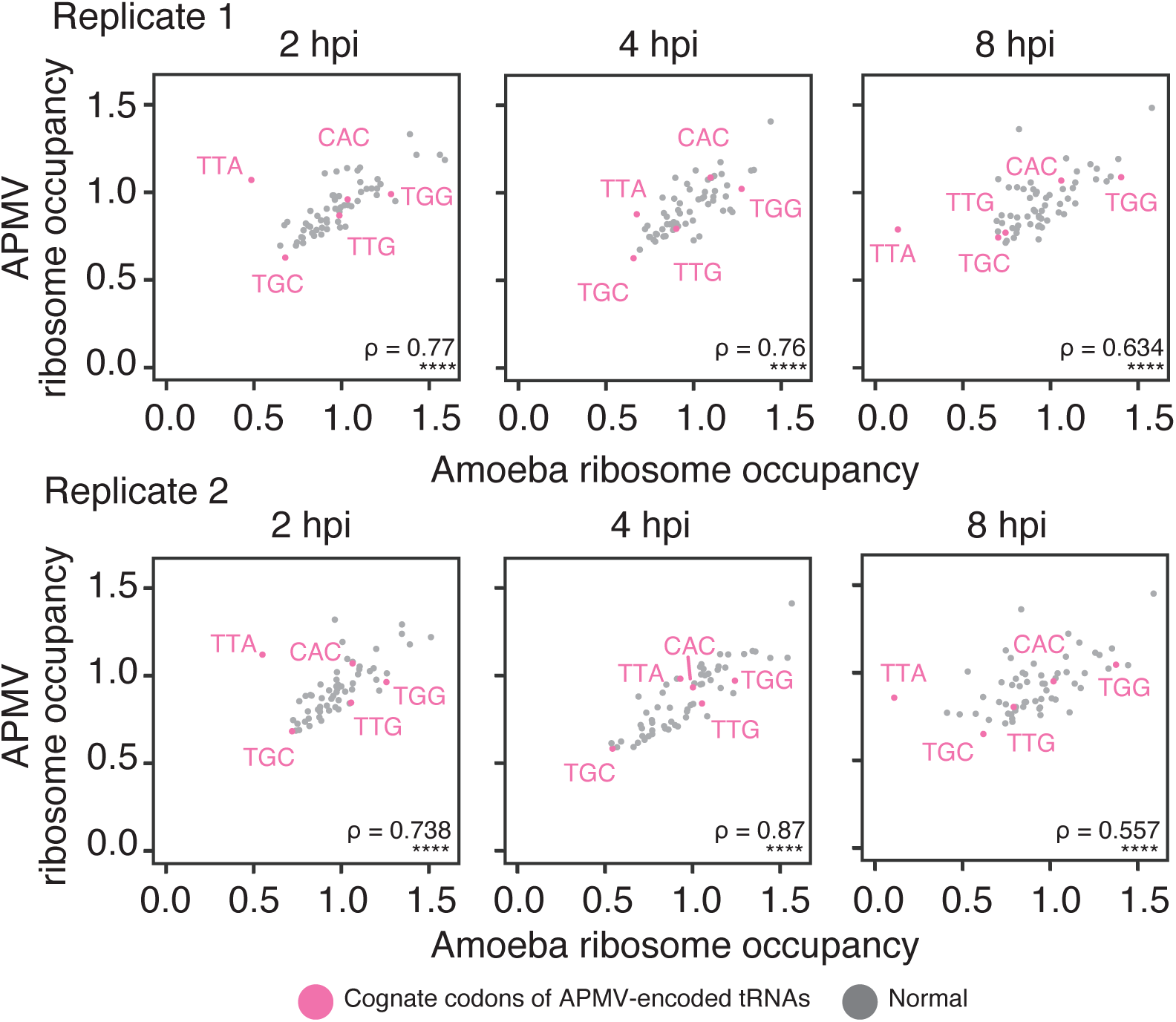
Correlation between amoeba and APMV ribosome occupancy. Scatter plots of the codon-specific ribosome occupancy on amoeba (horizontal axis) and APMV (vertical axis) mRNAs at different infection stages. Cognate codons of APMV-encoded tRNAs were coloured in pink. hpi, hours post-infection; ρ, Spearman’s correlation coefficients (two-tailed); ****, *p* <0.0001.

**Extended Data Table 1.** Average tRNA supply (*W* score) for AU3 and GC3 codons in each sample and *p* value from Wilcoxon rank sum test (one-side)

|  | Codon type |  | *p* value |  |
| --- | --- | --- | --- | --- |
| Sample | AU3 | GC3 | AU3 < GC3 | AU3 > GC3 |
| R1 0 hpi | 2.1671 | 2.8443 | 0.0587 | 0.9429 |
| R2 0 hpi | 2.1429 | 2.8509 | 0.0450 | 0.9563 |
| R1 2 hpi | 2.1709 | 2.8497 | 0.0587 | 0.9429 |
| R2 2 hpi | 2.1463 | 2.8408 | 0.0457 | 0.9557 |
| R1 4 hpi | 2.1051 | 2.8212 | 0.0587 | 0.9429 |
| R2 4 hpi | 2.0956 | 2.8264 | 0.0464 | 0.9550 |
| R1 8 hpi | 2.1713 | 2.8126 | 0.0696 | 0.9323 |
| R2 8 hpi | 2.1598 | 2.8240 | 0.0580 | 0.9437 |

**Extended Data Table 2.** Average balance score for AU3 and GC3 codons in each sample and *p* value from Wilcoxon rank sum test (one-side)

|  | Codon type |  | *p* value |  |
| --- | --- | --- | --- | --- |
| Sample | AU3 | GC3 | AU3 < GC3 | AU3 > GC3 |
| R1 0 hpi | 0.3858 | 0.0991 | 1.0000 | 1.2017e-06 |
| R2 0 hpi | 0.3656 | 0.0979 | 1.0000 | 1.2017e-06 |
| R1 2 hpi | 0.3024 | 0.3227 | 2.5347e-01 | 7.5113e-01 |
| R2 2 hpi | 0.2568 | 0.3004 | 1.8112e-01 | 8.2265e-01 |
| R1 4 hpi | 0.2116 | 0.3451 | 9.1690e-03 | 9.9118e-01 |
| R2 4 hpi | 0.1773 | 0.3326 | 3.7242e-03 | 9.9643e-01 |
| R1 8 hpi | 0.1445 | 0.3237 | 5.2970e-04 | 9.9950e-01 |
| R2 8 hpi | 0.0796 | 0.3099 | 1.5285e-06 | 1.0000 |

**Extended Data Table 3.** A-site offset for ribosome footprints of amoeba and APMV mRNAs.

| Amoeba |  | APMV |  |
| --- | --- | --- | --- |
| Length | A-site offset | Length | A-site offset |
|  |  | 18 | 12 |
| 19 | 15 | 19 | 13 |
| 20 | 15 | 20 | 14 |
| 21 | 15 | 21 | 15 |
| 22 | 16 | 22 | 16 |
| 23 | 16 | 23 | 9 |
| 25 | 11 | 24 | 10 |
| 26 | 12 | 25 | 11 |
| 27 | 15 | 26 | 12 |
| 28 | 15 | 27 | 13 |
| 29 | 15 | 28 | 14 |
| 30 | 15 | 29 | 15 |
| 31 | 16 | 30 | 15 |
| 32 | 16 | 31 | 16 |
|  |  | 32 | 16 |

